# The cell cycle oscillator and spindle length set the speed of chromosome separation in *Drosophila* embryos

**DOI:** 10.1101/2024.06.17.598879

**Authors:** Yitong Xu, Anna Chao, Melissa Rinaldin, Alison Kickuth, Jan Brugués, Stefano Di Talia

**Affiliations:** Department of Cell Biology, Duke University Medical Center, Durham NC 27705, USA; Duke Center for Quantitative Living Systems, Duke University, Durham NC 27710, USA; Cluster of Excellence Physics of Life, TU Dresden, Dresden, 01307 Germany; Max Planck Institute of Molecular Cell Biology and Genetics, Dresden, 01307 Germany; Center of Systems Biology, Dresden, 01307 Germany

## Abstract

Anaphase is tightly controlled in space and time to ensure proper separation of chromosomes. The mitotic spindle, the self-organized microtubule structure driving chromosome segregation, scales in size with the available cytoplasm. Yet, the relationship between spindle size and chromosome movement remains poorly understood. Here, we address how the movement of chromosomes changes during the cleavage divisions of the *Drosophila* blastoderm. We show that the speed of chromosome separation gradually decreases during the 4 nuclear divisions of the blastoderm. This reduction in speed is accompanied by a similar reduction in the length of the spindle, thus ensuring that these two quantities are tightly linked. Using a combination of genetic and quantitative imaging approaches, we find that two processes contribute to controlling the speed at which chromosomes move at mitotic exit: the activity of molecular motors important for microtubule depolymerization and sliding, and the cell cycle oscillator. Specifically, we found that the levels of Klp10A, Klp67A, and Klp59C, three kinesin-like proteins important for microtubule depolymerization, and the level of microtubule sliding motor Klp61F (kinesin-5) contribute to setting the speed of chromosome separation. This observation is supported by quantification of microtubule dynamics indicating that poleward flux rate scales with the length of the spindle. Perturbations of the cell cycle oscillator using heterozygous mutants of mitotic kinases and phosphatases revealed that the duration of anaphase increases during the blastoderm cycles and is the major regulator of chromosome velocity. Thus, our work suggests a potential link between the biochemical rate of mitotic exit and the forces exerted by the spindle. Collectively, we propose that the cell cycle oscillator and spindle length set the speed of chromosome separation in anaphase.

## Results

In *Drosophila* embryos, early development is characterized by rapid and synchronous syncytial nuclear divisions [1, 2]. At the blastoderm stage, multiple nuclear divisions occur simultaneously on the surface of the embryo. These mitoses drive a reduction in the spacing among nuclei, which, in turn, results in smaller mitotic spindles. To elucidate the relationship between spindle size and chromosome separation, we used confocal live imaging to study anaphase in early fly embryos.

Using embryos maternally expressing histone tagged with GFP (His2Av-GFP) and a microtubule fluorescent reporter (mCherry fused to the Tau microtubule-binding domain) [3], we imaged nuclear divisions and microtubule dynamics from cycle 10 to cycle 13 (Fig 1A-B). As the nuclear cycles progressed, the number of nuclei at the surface of the embryo increased exponentially, while the spacing among nuclei proportionally reduced. This means chromosomes in an early cycle (e.g. cycle 10) possess larger room for separation than chromosomes in a later cycle (e.g. cycle 13). Likewise, mitotic spindles were proportionally smaller as the embryo approached later cycles (Fig 1A). We used the histone signal to segment chromosomes and tracked their segregation in individual nuclei [4]. To quantify the velocity of chromosome separation, we measured the speed at which the leading edges of segregating sister chromatids move apart from each other (Fig 1C, S1). These measurements revealed a compelling finding: the speed at which chromosomes separate during anaphase exhibits a scaling relationship with spindle length (estimated here as the maximum distance of chromosome separation, see Fig S1) (Fig 1H). This scaling ensures that chromosome separation has a similar duration-approximately 70 seconds- in all nuclear cycles (Fig 1C). Moreover, the dynamics of the distance between sister chromosomes can be collapsed for all cycles when normalized by total distance that is covered in each cycle (Fig 1D), suggesting that the dynamics are essentially indistinguishable when rescaled for spindle length. Additionally, we found that both the movement of the chromosome towards the spindle pole (Anaphase A) and the movement of spindle poles away from each other (Anaphase B) demonstrated scaling with spindle length (Fig 1E-G). Quantitative comparison of the chromosomes movements due to these two processes confirmed that, as expected, chromosome separation is dominated by Anaphase A in fly embryos [5, 6] (Fig 1F-G, S1). Collectively, these observations point to an interesting correlation between the speed of chromosome separation and the length of the mitotic spindle.

**Figure 1.**
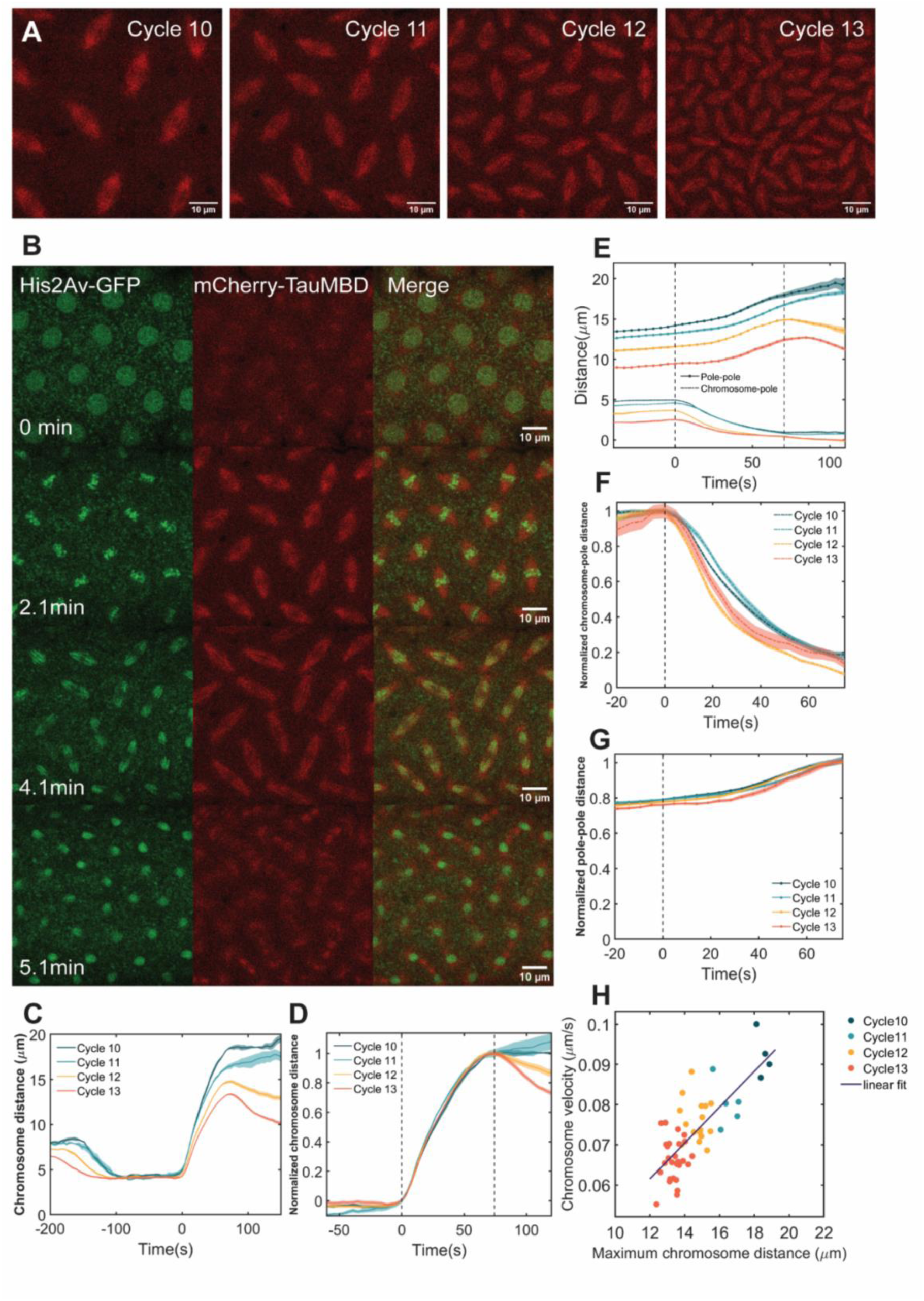
Chromosome velocity during anaphase scales with spindle length. (A) Mitotic spindles in the *Drosophila* embryo from syncytial cycle 10 to 13, labeled with mCherry-Tau microtubule binding domain. (B) Chromosome and microtubule dynamics during mitosis in a cycle 11 embryo. From top to bottom: prophase, metaphase, anaphase, telophase. (C) Distance between the leading edges of sister chromosomes during anaphase as a function of time. (D) Normalized chromosome distance from (C) by the maximum distance. Left dotted line: anaphase onset. Right dotted line: end of chromosome movement during anaphase, when chromosome distance reached its maximum or plateau. (E) Pole-to-pole distance and chromosome-to-pole distance in a His-RFP γ-Tubulin-GFP embryo during anaphase from cycle 10 to 13. (F and G) Both chromosome-to-pole distance and pole-to-pole distance could be rescaled across cycles. (H) The average chromosome velocity during anaphase scales with the maximum chromosome distance, which serves as a proxy of spindle length.

To elucidate the connection between spindle length and chromosome separation speed, we first considered the potential role of microtubule dynamics. To this end, we conducted a series of experiments to quantify different parameters of spindle microtubule dynamics. First, we investigated microtubule density along the pole-to-pole axis during mitosis. As chromosomes separate and spindles elongate, the spatial distribution of microtubules remains largely unchanged, exhibiting no discernable correlation with chromosome (and kinetochore microtubules) position, until the spindle disassembles (Fig 2A-B). Thus, in early anaphase, kinetochore microtubules likely represent a small fraction of spindle microtubules. The decrease in spindle length from cycle 10 to cycle 13 is accompanied by a decrease in microtubule density (Fig 2C-D). Secondly, we performed tracking of microtubule plus-ends and quantified the rate of microtubule polymerization, using embryos expressing microtubule plus-end binding protein EB1-GFP [7–10]. Our analysis revealed that the polymerization velocity of microtubules has a significant, although slight, dependency on spindle length, as polymerization velocity increases about 20% when spindle length doubles (Fig 2E-F). A similar positive correlation between spindle length and microtubule polymerization velocity has been observed in zebrafish, *C. elegans* and sea urchin [8, 11]. However, the extent of this correlation quantitatively changes in these organisms: a strong dependency of polymerization speed on spindle length is observed in sea urchin and *C. elegans*, while a small dependency is observed in zebrafish, similar to the one seen here for *Drosophila*.

**Figure 2.**
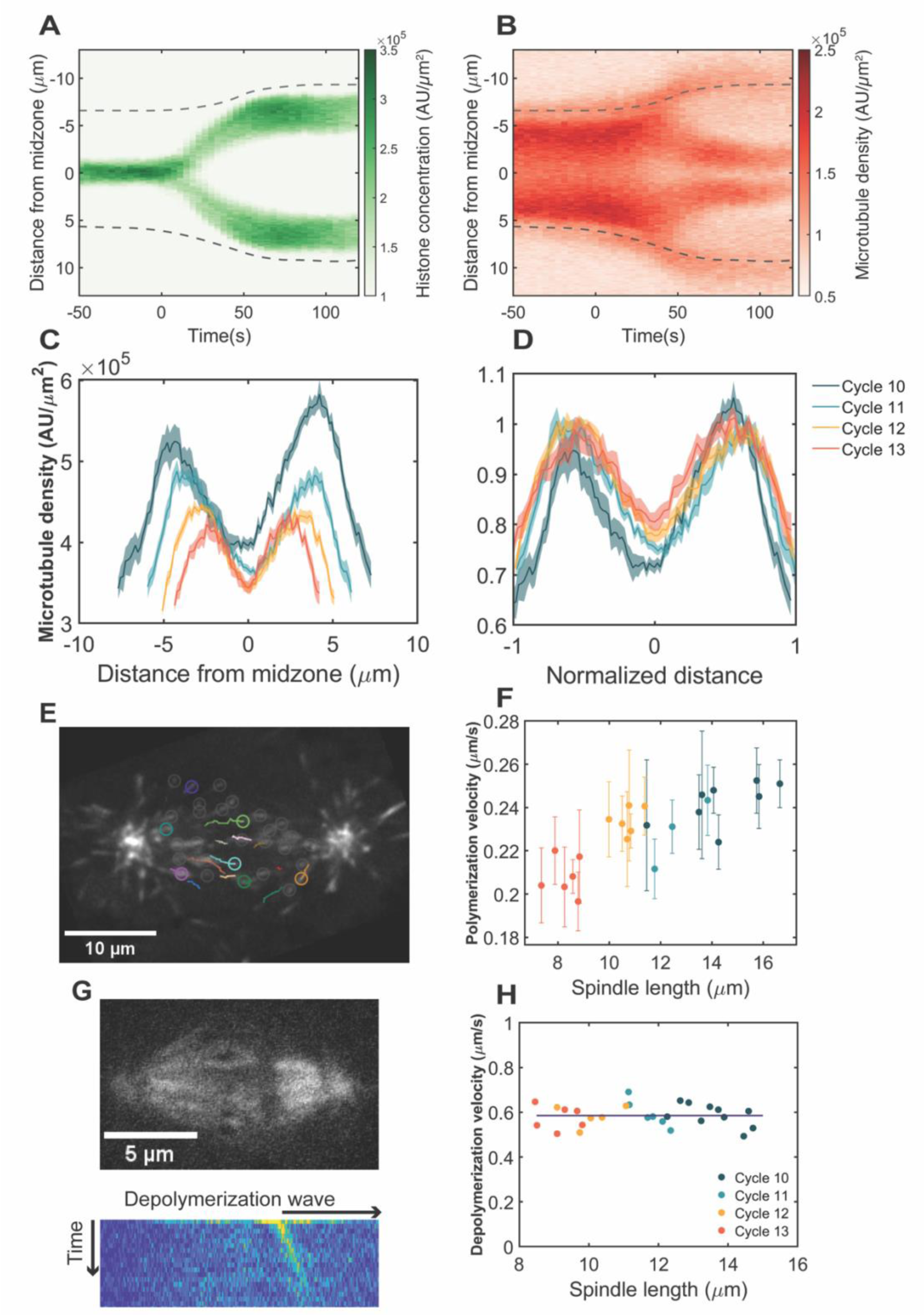
Microtubule dynamics contribute to spindle scaling in the *Drosophila* embryo. (A and B) Kymographs of chromosome separation (A) and spindle dynamics (B) in a cycle 11 embryo, labeled with His2Av-GFP and mCherry-Tau microtubule binding domain. T = 0 indicates anaphase onset. Dashed lines indicate approximated centrosome positions. (C) Line scans of microtubule density along the spindle long axis during metaphase in one embryo (n= 5 spindles for each cycle, mean± SEM). (D) Rescaled microtubule density from cycle 10 to 13. (E) Microtubule plus end tracking with EB1-GFP in a cycle 12 embryo. (F) Microtubule polymerization velocity as a function of spindle length. Each data point represents the mean velocity ± SEM of all tracked comets in one spindle. (G) Laser ablation of microtubules in a cycle 11 embryo, labeled with Jupiter-GFP. A wave of depolymerization was visualized after projecting the differential intensity onto the spindle long axis. (H) Microtubule depolymerization velocity as a function of spindle length.

This observation argues that microtubule polymerization contributes partially to the modulation of spindle length in the *Drosophila* blastoderm. Finally, we employed femto-second laser ablation to sever microtubules within the metaphase spindle, inducing microtubule depolymerization [12, 13]. A consistent rate of depolymerization of approximately 0.6µm/s (35µm/min) was observed, regardless of the specific cell cycle stage or spindle length, which demonstrated that the rate of depolymerization of unstable microtubules does not change during cycle 10 to 13 (Fig 2G-H). This observation and the fact that the measured value is consistent with values observed in other systems argue that this constant rate of depolymerization is set by the intrinsic properties of microtubule catastrophe dynamics. These observations on microtubule dynamics are similar to previous findings in zebrafish [8], suggesting a conserved mechanism for the scaling of spindle size during *Drosophila* blastoderm divisions. However, they do not explain the relationship between spindle length and chromosome speed.

Poleward movement of chromosomes (Anaphase A) is achieved by the shortening of kinetochore-associated microtubules, while the separation of opposite spindle poles (Anaphase B) is driven by the sliding of interpolar microtubules. Both processes involve forces generated on the microtubules by motor proteins. Thus, we turned our attention to the molecular motors involved in those processes and more specifically motors that play a role in shortening kinetochore-associated microtubules, given the dominant contribution of Anaphase A to chromosome separation. To this end, we first analyzed microtubule poleward flux, that is the continuous movement of tubulin subunits towards the centrosome [14–16]. We note that for kinetochore-associated microtubules during anaphase, poleward flux is driven by microtubule depolymerization at both the centrosome (minus end) and at the kinetochore (plus end), due to the activity of specific motors [17, 18]. For polar microtubules, poleward flux could be due to either depolymerization at the centrosome or microtubule sliding. To estimate poleward flux in the fly embryos, we employed a transgenic line in which tubulin is tagged with a photo-convertible tdEOS fluorescent protein that can be converted from green to red upon UV illumination [19]. To describe poleward fluxes in anaphase, we monitored spindle morphology under the confocal microscope in living embryos [20]. As nuclei approached anaphase onset, we photo-converted a small region of microtubules near the mid-spindle and tracked it for 20-30 seconds during early anaphase (Fig 3A). The calculated poleward flux rates were comparable to the speed of chromosome movement at anaphase onset and similar to previously reported values [16]. The flux rate showed a clear dependency on spindle length, suggesting that it might be implicated in setting the speed of chromosome movement in anaphase (Fig 3B).

**Figure 3.**
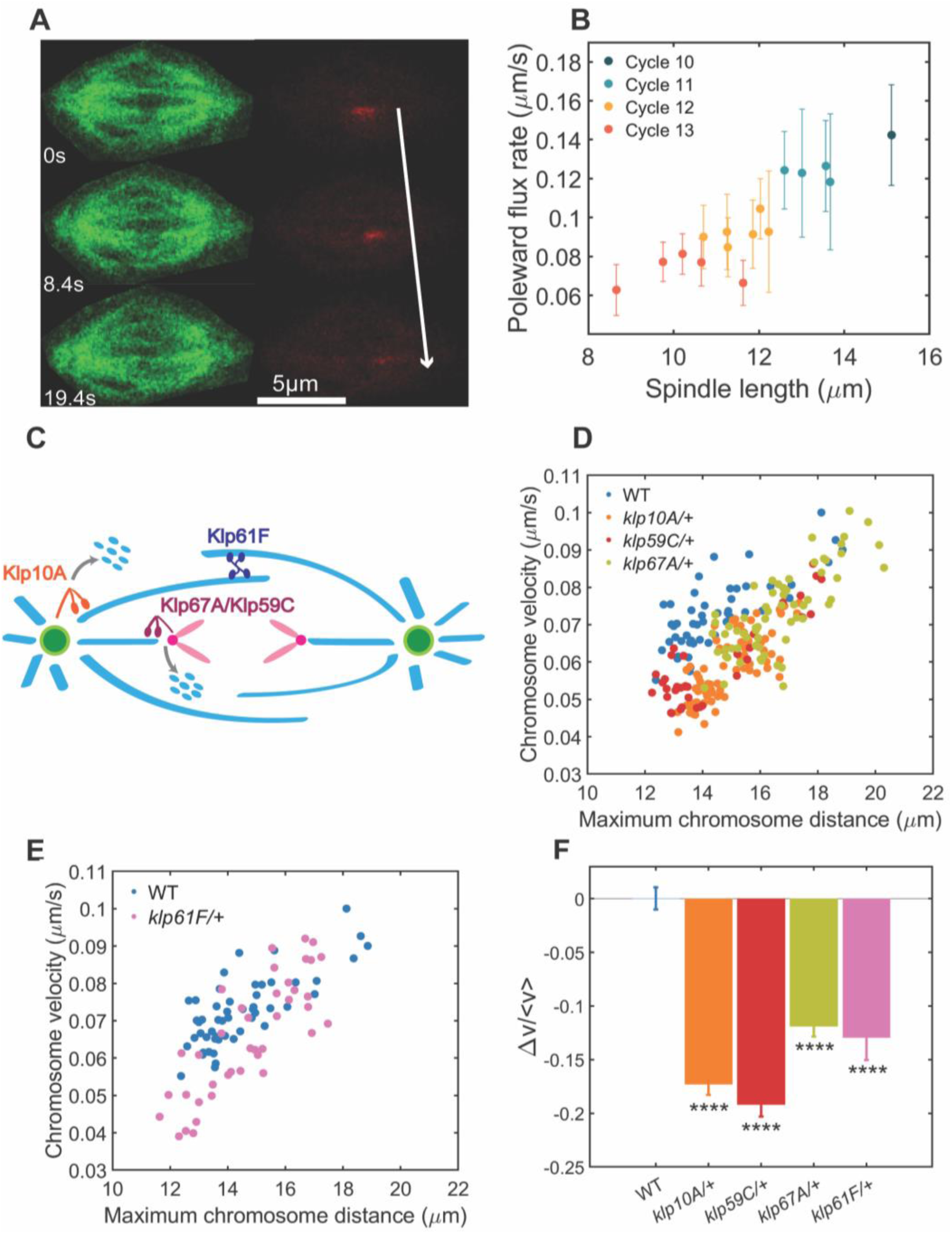
Microtubule-depolymerizing motors regulate chromosome velocity. (A) Photo-conversion of the microtubules using α-tubulin-tdEOS embryos. (B) Poleward flux rate scales with spindle length. (C) Microtubule-depolymerizing motors Klp10A, Klp59C and Klp67A contribute to chromosome-to-pole movement (Anaphase A), while microtubule-sliding motor Klp61F contributes to pole-pole separation (Anaphase B). (D) Correlation between chromosome velocity and maximum chromosome distance in one WT (n= 52), one Klp10A (n= 76) and one Klp67A (n= 65) heterozygous motor mutant. (E) Correlation between chromosome velocity and maximum chromosome distance in one WT (n= 52) and two Klp61F (n= 40) heterozygous motor mutants. (F) Average relative change in chromosome velocity between wild-type and heterozygous motor mutants (mean± SEM, **** P<0.0001).

To strengthen this point, we analyzed whether motors implicated in Anaphase A and B movement, namely the kinesin-13 motors, Klp10A and Klp59C, the kinesin-8 Klp67A, and the kinesin-5 Klp61F, are rate-limiting for chromosome movements (Fig 3C). We used heterozygous mutants to lower their activity without fully abrogating it. We analyzed chromosome velocity and spindle length in these mutants and, after correcting for changes in spindle length due to loss of motor function, found that they retain a strong relationship between chromosome speed and spindle length (Fig 3D). Notably, all heterozygous mutants display a significant reduction in the speed of chromosome separation for a given spindle length (Fig 3E). Similar effects were observed for motors acting at the centrosome (Klp10A) and at the kinetochore (Klp67A and Klp59C), suggesting that both processes contribute to chromosome movements in comparable manner (Fig 3E). Klp10A also localizes to the kinetochore, but its activity at the kinetochore is believed to be weaker than at the centrosome [18]. Perturbing the level of Klp61F, the microtubule sliding motor driving Anaphase B, also caused a slowdown of chromosome separation, as well as a small decrease in spindle length (Fig. 3E), concordant with previous data [21]. In addition to Anaphase B, Kinesin-5 can influence Anaphase A by coupling sliding interpolar MTs to kinetochore fibers, an idea further supported by previous more severe inhibition of Klp61F function by antibody-induced dissociation of the motor from spindles [21]. Collectively, these results suggest that the scaling of chromosome movement with spindle length is the result of changes in the rate of microtubule polymerization, depolymerization and sliding. Moreover, our genetic experiments implicate multiple motors in this process, thus suggesting that there might be a global mechanism controlling the activity of molecular motors in anaphase.

A natural candidate for such regulation is the cell cycle oscillator, as the activity of the motors must be controlled in space and time during the cell cycle. Specifically, we hypothesized that the rates of phosphorylation and dephosphorylation of mitotic targets involved in the function of the spindle might influence the speed of chromosome separation. Changes in these rates likely set the rate of completion of anaphase, which in turn could contribute to setting the speed of chromosome movement by influencing the activity of mitotic targets, such as molecular motors, involved in spindle function. To measure anaphase rate in different cell cycles, we defined anaphase duration, as the time period from the initiation of chromosome segregation to nuclear envelope reformation (Fig 4A-C), which we operationally used as the hallmark event to score completion of anaphase [22, 23] (Fig 4A-B). We used the intensity of nuclear-localized GFP to estimate the time when nuclear envelope integrity was reestablished. Imaging this probe together with histones showed that the reformation of nuclear envelope starts after the segregation of chromosomes is completed (Fig S3). This analysis showed that the duration of anaphase gets progressively longer from nuclear cycle 10 to 13 and, most importantly, that there is a strong correlation between anaphase duration and the speed of chromosome separation (Fig 4D).

**Figure 4.**
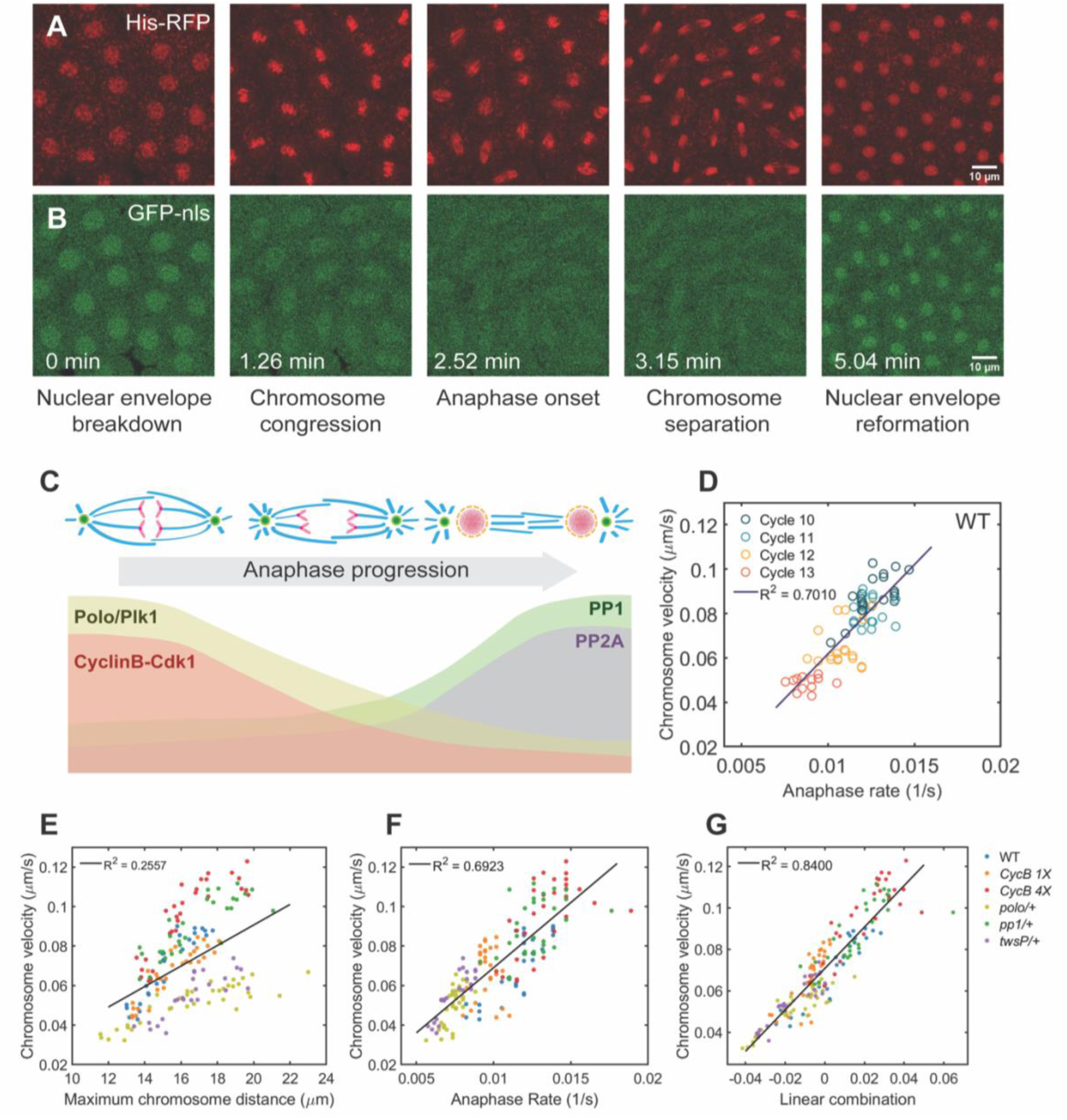
The cell cycle oscillator and spindle length together set the speed of chromosome movement in anaphase. (A and B) Dynamics of chromosome and nuclear localization signals (NLS) in a cycle 11 embryo, from nuclear envelope breakdown to nuclear envelope reformation. (C) Schematic of anaphase progression and the corresponding activity of mitotic kinases and phosphatases. (D) Chromosome velocity scales with anaphase rate in wild-type embryos. (E) Chromosome velocity as a function of maximum chromosome distance in heterozygous cell cycle mutants. (F) Chromosome velocity as a function of anaphase rate in heterozygous cell cycle mutants. (G) Chromosome velocity scales with a linear combination of anaphase rate (70%) and spindle length (30%).

To gain further insight on this correlation, we investigated the relationship between chromosome speed and spindle length in embryos heterozygous for several regulators of the cell cycle (Fig 4E). Remarkably, we found that these mutations disrupted the scaling relationship. Embryos with one less copy of Cyclin B (1*x cycB*) displayed slightly slower chromosome velocity. On the contrary, embryos with two extra copies of Cyclin B (*4x cycB*) exhibited a faster chromosome separation speed. These observations suggest that Cyclin B-Cdk1 plays a role in setting the speed of chromosome separation. We also observed a reduction of chromosome velocity in *polo* (Plk1) and *twsP* (the B55 regulatory subunit of the PP2A phosphatase) heterozygous mutants. In *polo/+* embryos, the nuclei migrating from the inner regions of the embryo reached the cortex at cycle 9 [24], that is one cycle earlier than in other genotypes, resulting in larger spindles at the onset of the blastoderm stage. Moreover, some *polo/+* embryos failed to finish cycle 13, due to cell cycle defects and excessive crowding of the nuclei at the embryo cortex. The *twsP/+* embryo displayed normal spindle length despite slower chromosome speed than wild-type. Finally, we found that *PP1-87B* and *PP1-96A* double heterozygous mutant embryos had slightly larger spindles and speeds than wild-type (similar to *4x cycB*) suggesting that Cdk1 and PP1 might have opposite impacts on chromosome separation. Collectively, these results revealed a major role for the components of the cell cycle oscillator in setting the speed of chromosome separation.

The cell cycle oscillator coordinates mitotic events in space and time (Fig 4C). In mitosis, Cdk1 activity represses cytoplasmic microtubule polymerization by affecting several microtubule-associated proteins (MAPs) and promotes spindle assembly and chromosome alignment [25]. Polo kinase also controls several aspects of spindle and centrosome behaviors [26–28]. The decrease of Cdk1 activity triggers the onset of anaphase and the migration of chromosomes towards the poles. As Cdk1 and Polo activities decrease in mitosis, mitotic substrates are dephosphorylated mainly by PP1/PP2A phosphatases [29, 30]. Together the balance of mitotic kinases and phosphatases controls all events at mitotic exit, including chromosome decondensation and the reformation of the nuclear envelope. Analysis of the duration of anaphase indicated that such duration is longer in the *cycB/+*, *polo/+* and *twsP/+* heterozygous embryos (Fig 4F). Conversely, *4x cycB* embryos and PP1 heterozygous embryos showed shorter anaphase durations than wild-type. These findings support the hypothesis that the activity of cell cycle regulators set the rate of completion of anaphase.

Given the observed changes in anaphase duration, we tested whether the speed of chromosome separation could be explained by these changes. In support of this idea, we found a strong correlation between chromosome velocity and anaphase rate, and most importantly, that such correlation holds essentially for all the mutants, so that all the data can be collapsed on a single relationship (Fig 4F). This observation suggests that the rate of progression through anaphase is a strong predictor, and most likely a major regulator, of the speed of chromosome separation. Careful inspection of the data showed that in regions where the anaphase rate overlapped between wild-type and mutant embryos, the embryos with larger spindles tended to have slightly higher speeds. This observation suggested that there is still some contribution to the speed of chromosome movement from processes independent of anaphase duration. We speculated that this residual contribution arises from differences in spindle length and/or microtubule dynamics [31]. Thus, we used a simple linear model to test how much of the data can be captured by a linear combination of anaphase duration and spindle length. After centering and normalization of the two variables, we found that a linear combination of anaphase rate and spindle length can predict chromosome speed with high accuracy in all the mutants (R^2^=0.84). This analysis suggests that anaphase rate accounts for 70% of the dependency of chromosome velocity whereas residual contributions from spindle length (or a closely correlated variable) might explain the remaining 30%. Collectively, these results argue that the speed of chromosome separation is set by a combination of biochemical cues from the cell cycle oscillator and mechanical cues from the mitotic spindle.

## Discussion

Anaphase is the culmination of mitosis, when duplicated chromosomes segregate and move towards opposite poles. In many cells, anaphase accounts for a very short portion of the cell cycle [32–34]. The rapidity and precise spatial control needed for a successful anaphase could pose challenges for the accuracy of chromosome segregation. Here, we carefully characterized the movement of chromosomes during anaphase as *Drosophila* embryos proceed through the blastoderm cycles. We found that the average speed of chromosome separation during these cycles is in large part controlled by the rate at which nuclei complete anaphase. Spindle length (and/or microtubule dynamics) and the concentration of molecular motors involved in microtubule depolymerization further contribute to this process ensuring a tight relationship between chromosome separation and spindle length in *Drosophila* blastoderm embryos.

The mechanisms of chromosome segregation during anaphase vary across different biological systems. In *Drosophila* embryos and human cell lines, chromosome to pole movement (Anaphase A) dominates the total chromosome movement. On the contrary, in *C. elegans* embryos, chromosome movement is almost solely achieved by pole-pole separation (Anaphase B) [6]. Our analysis of microtubule dynamics and our genetic manipulations of molecular motors argue the speed of chromosome separation is mainly set by the activity of microtubule depolymerizing motors. Notably, all three microtubule depolymerizing motors, Klp10 A, Klp67A and Klp59C, as well as kinesin-5 Klp61F, contribute to setting chromosome speed in anaphase. Two major drivers of Anaphase A movement have been proposed: microtubule depolymerization at the centrosome and the kinetochore. It has been debated which of the two mechanisms contributed more.

Our analysis of poleward fluxes and of mutants reducing the activity of molecular motors argues that both processes contribute to a comparable extent to Anaphase A, which might explain why it has been difficult previously to conclusively establish their relative importance. We note that the microtubule poleward flux rate we observed in this work is comparable to results in Maddox et al [16].

Our results support a model in which the rate at which anaphase is completed is a major determinant of the speed of chromosome movement. This observation can be linked to the role of molecular motors by proposing that a major function of the cell cycle oscillator is to set the activity of Klp10A, Klp67A and Klp59C and thus control the speed of Anaphase A movement. This model is supported by previous experiments showing that phosphorylation by mitotic kinases can influence the activity of MCAK, the major microtubule depolymerizing kinesin in human cells [35]. Notably, we found that the rate of anaphase completion depends similarly on the activity of mitotic kinases Polo and CycB-Cdk1 and the phosphatase PP2A. Genetic manipulations that decrease the activity of all the three enzymes result in slower progression through anaphase. Our analysis also revealed that PP2A-B55, rather than PP1, is the rate-limiting phosphatase for completion of anaphase, or at least for timing nuclear envelope reformation, consistent with previous findings [23]. These observations suggest that the rate of completion of anaphase is likely to depend on the feedback mechanisms that drive both phosphorylation and dephosphorylation of mitotic targets, as well as the feedback mechanisms by which Polo and Cdk1 control PP2A and vice versa. Understanding how these feedback mechanisms operate to control the dynamics of anaphase will reveal important new insights on mitotic regulation.

Our experiments also show that cell cycle dynamics cannot fully explain the speed of chromosome separation and that other molecular processes that correlate with or are controlled by spindle length must be involved. A possibility is that the number of molecular motors available is titrated out by the increasing number of kinetochores and centrosomes associated with nuclear divisions. Consistent with this, using a Klp10A-GFP transgenic line, we observed a slight decrease in the Klp10A concentration at centrosomes from cycle 11 to cycle 13 (Fig S2). Alternatively, the effects of spindle length on chromosome separation could arise from geometric or physical effects, for example via length-dependent processes if microtubule length were to scale with spindle size or other mechanisms by which forces might scale with spindle length [36]. These possible mechanisms are currently unclear and remain to be identified.

Scaling of spindle size with available cytoplasm has been widely reported both *in vivo* and *in vitro*, and it is believed that microtubule polymerization and nucleation set spindle scaling in systems of all sizes [8, 37]. In this study, we revealed an association between spindle scaling and its cellular function, that is chromosome segregation. Understanding if this association can be generalized to other systems, in particular to other embryos undergoing reductive cleavage divisions, could reveal a conserved link between these two fundamental subcellular processes. Furthermore, understanding if and how the scaling identified here is influenced by the syncytial and multi-nucleated nature of the cytoplasm will be important. In the future, developing an integrated model that combines the dynamics of phosphorylation levels, microtubule quantity and dynamics, and cell cycle progression will be essential to elucidate the mechanism of scaling of chromosome speed.

## Supplemental information

Figures S1-3

Videos S1: Nuclear division and spindle dynamics in the Drosophila blastoderm.

Video S2: Growing microtubule plus-ends labeled with EB1-GFP.

Video S3: Laser ablation of a mitotic spindle.

Video S4: Photo-conversion of kinetochore-associated microtubules.

## Acknowledgements

We thank the Bloomington Drosophila Stock Center, Sharyn Endow, David Glover, Michael Goldberg and Pavel Tomancak for providing stocks. We thank Mary Elting, Sharyn Endow, Christine Field and Tim Mitchison for discussions. We thank members of the Di Talia lab for discussions. This work was supported by NIH R01-GM122936 and R35-GM153490. M.R. acknowledges funding from the Human Frontier of Science (Postdoctoral cross disciplinary fellowship LT000920/2020-C) and the European Molecular Biology Organization (Postdoctoral fellowship EMBO ALTF 597-2021). J.B., M.R., and A.K. acknowledge support from the Deutsche Forschungsgemeinschaft (DFG, German Research Foundation) under Germany’s Excellence Strategy – EXC-2068– 390729961-Cluster of Excellence Physics of Life of TU Dresden.

## Author contribution

Conceptualization, Y.X. and S.D.; Methodology, Y.X., J.B. and S.D.; Software, Y.X.; Formal Analysis, Y.X. and S.D.; Investigation, Y.X., A.C., M.R. and A.K.; Writing, Y.X. and S.D.; Visualization, Y.X. and S.D.; Supervision, S.D. and J.B., Funding Acquisition, S.D. and J.B.

## RESOURCE AVAILABILITY

### Materials availability

This study did not generate new fly lines or reagents.

### Data and code availability

All the microscopy data reported in this paper will be shared by the lead contact upon request.

All original code has been deposited at Github at the following link and is publicly available as of the date of publication: https://github.com/Yitong-Xu/Scaling2024 Any additional information required to reanalyze the data reported in this paper is available from the lead contact upon request.

## EXPERIMENTAL MODEL AND STUDY PARTICIPANT DETAILS

### Fly lines and husbandry

Klp10A P mutant flies were raised at room temperature (∼21°C) on glucose food (Archon, Cat #D210). All other flies were raised at room temperature on standard molasses food (Archon, Cat #B20101).

## METHOD DETAILS

### Embryo collection and processing

Before imaging, adult flies with genotypes of interest were housed in a cage covering an apple juice plate at 25°C, supplemented with yeast paste. Embryos were collected over 2 hours on a fresh plate, dechorionated with 50% bleach for 1 minute, and mounted in Halocarbon oil 27 on a gas-permeable membrane with coverslips.

### Microscopy

Embryos were imaged on a Leica SP8 laser scanning confocal microscope equipped with a Leica 20x oil-immersion objective 0.75NA (HC PL APO CS2 20x/0.75 IMM) unless otherwise noted.

### Microtubule plus end imaging

Embryos were imaged with a spinning disk confocal microscope (IX83 Olympus microscope with CSU-X1 Yokogawa disk) connected with two iXon DU-897 back-illuminated EMCCD camera (Andor). Experiments were acquired using an Olympus 100x silicon oil objective1.35 NA and imaged at 1-2 frames per second.

### Laser ablation experiments

With the Jupiter-GFP line, metaphase spindles were imaged using a spinning disk confocal microscope (Nikon Ti Eclipse, Yokogawa CSU-X1) equipped with an EMCCD camera (iXon DU-888 or DU-897, Andor) and a 100x oil-immersion objective. Images were acquired with the Andor iQ software. Laser ablation was performed according to Rieckhoff et al on a custom-built femtosecond laser microsurgery system. Briefly, line cuts parallel to the spindle equator were induced by moving the sample with a high-precision piezo stage (PInano) relative to the stationary cutting laser. The ablation was controlled by a custom-written software managing the piezo-stage and a mechanical shutter in the optical path. Each embryo was cut only once and imaged at intervals of 200-300ms/frame.

### Photo-conversion experiments

Photo-conversion experiments were performed on the Leica SP8 microscope with the FRAP Module in the Leica Application Suite X (LAS X). Experiments were acquired using a Leica 63x oil objective 1.40 NA (HC PL APO CS2 63x/1.40 OIL). Spindle microtubules were excited with 0.1% 405nm laser for 1 millisecond to induce photo-conversion from GFP to RFP.

## QUANTIFICATION AND STATISTICAL ANALYSIS

### Chromosome segmentation and tracking

Movies of the histone channel were loaded in ilastik for chromosome segmentation using the pixel classification workflow. Pixels were manually annotated as either chromosome or cytosol to train the classifier. Training was considered complete when the live output aligned with visual judgement. A prediction map for chromosome segmentation was generated. The raw movie and prediction map were reloaded into ilastik for the tracking or manual tracking workflow, with the division events manually labeled. For automatic tracking, the correctness of the tracking was manually verified before further quantification. Either the maximum number of trackable nuclear division events in frame or at least five divisions were recorded for each cycle.

### Quantification of chromosome distance and chromosome velocity

For segmented chromosome undergoing mitosis, a bounding box was drawn surrounding a single nucleus or sister chromosomes. The length of the bounding box along the division axis was quantified as chromosome distance in real-time. Anaphase onset was determined as the first frame when chromosome distance started to increase after metaphase. The total chromosome movement during anaphase was quantified from anaphase onset till the chromosome distance plateaued (e.g. in cycles 10 and 11) or reached maximum before recoil (e.g. in cycles 12 and 13). Average chromosome velocity was calculated as total chromosome movement divided by the duration of the movement. To compare the velocity among genotypes controlling for spindle length, data points were divided into 4 bins based on spindle lengths (ranging from 12 µm to 20 µm). The average chromosome velocity of the wild-type in these bins was quantified as a refence velocity <v_WT_>. For each bin, the relative change in velocity was quantified as (v_mutant_-<v_WT_>)/<v_WT_>. Relative changes in all bins were summarized for each genotype.

### Quantification of Anaphase A and B movement

Using the TrackMate plugin in FIJI, centrosomes marked with γTubulin were detected with the LoG detector and tracked with the LAP tracker, allowing for splitting. Centrosome tracks were manually curated and matched with the corresponding chromosome tracks. Anaphase B movement was quantified by measuring the separation of two centrosomes at opposite spindle poles. Anaphase A movement was calculated by subtracting Anaphase B movement from the total chromosome movement.

### Quantification of microtubule density

For spindles at metaphase, a line of defined thickness (2µm) along the spindle long axis was used to measure fluorescence intensity and calculate the density profile.

### Quantification of microtubule polymerization velocity

EB1 comets in the spindle region were tracked with the TrackMate plugin in the FIJI software, applying the LoG detector and simple LAP tracker. Tracks were filtered by duration (∼ 3-15s) and linearity (> ∼ 0.9) and then manually screened. The speeds of all correctly tracked comets within each spindle were averaged to represent the microtubule polymerization velocity for that spindle.

### Quantification of microtubule depolymerization velocity

The amount of depolymerized microtubule during a time interval was calculated by subtracting raw images with a time difference of 0.4∼0.6s from each other and integrating these differential intensities perpendicular to the spindle long axis. Depending on the position of the cut, the integrated differential intensities along the spindle long axis showed one or two well-defined peaks. The peaks moved toward the nearest pole following ablation. The more prominent peak was fit to a Gaussian function to quantify the position of the maximum. The position of the maxima over time was fit to a line to determine the microtubule depolymerization velocity.

### Quantification of poleward flux rate

To analyze the poleward flux, for images in the photo-converted channel, we computed the average fluorescence intensity along the spindle length and evaluated the points where this quantity crosses a value close to half-max, estimated as half of the 95 percentile of fluorescence intensity values. These points defined the ends of the photo-converted region. For instances when the spindles remain a constant length and where microtubules on both sides of the mid-spindle were properly labeled, the speed of poleward flux was estimated as half of the speed at which the two ends moved apart. For instances where microtubules were labeled only on one side of the spindle, we computed the speed as the distance between the photo-converted end and the closest centrosome. The positions of centrosomes were estimated by computing the initial and final positions where intensity in the green (non-converted) channel crossed a value close to half-max (half of the 95 percentile of fluorescence intensity values).

### Quantification of nuclear envelope reformation

The Histone-RFP channel in the time-lapse movie was used to segment nuclei and track nuclear division with ilastik. The nuclear concentration of GFP-NLS signal, together with chromosome distance, was plotted as a function of time. The onset of nuclear envelope reformation was inferred from the time point when the GFP-NLS concentration began to increase after chromosome separation.

---

**Figure S1.**
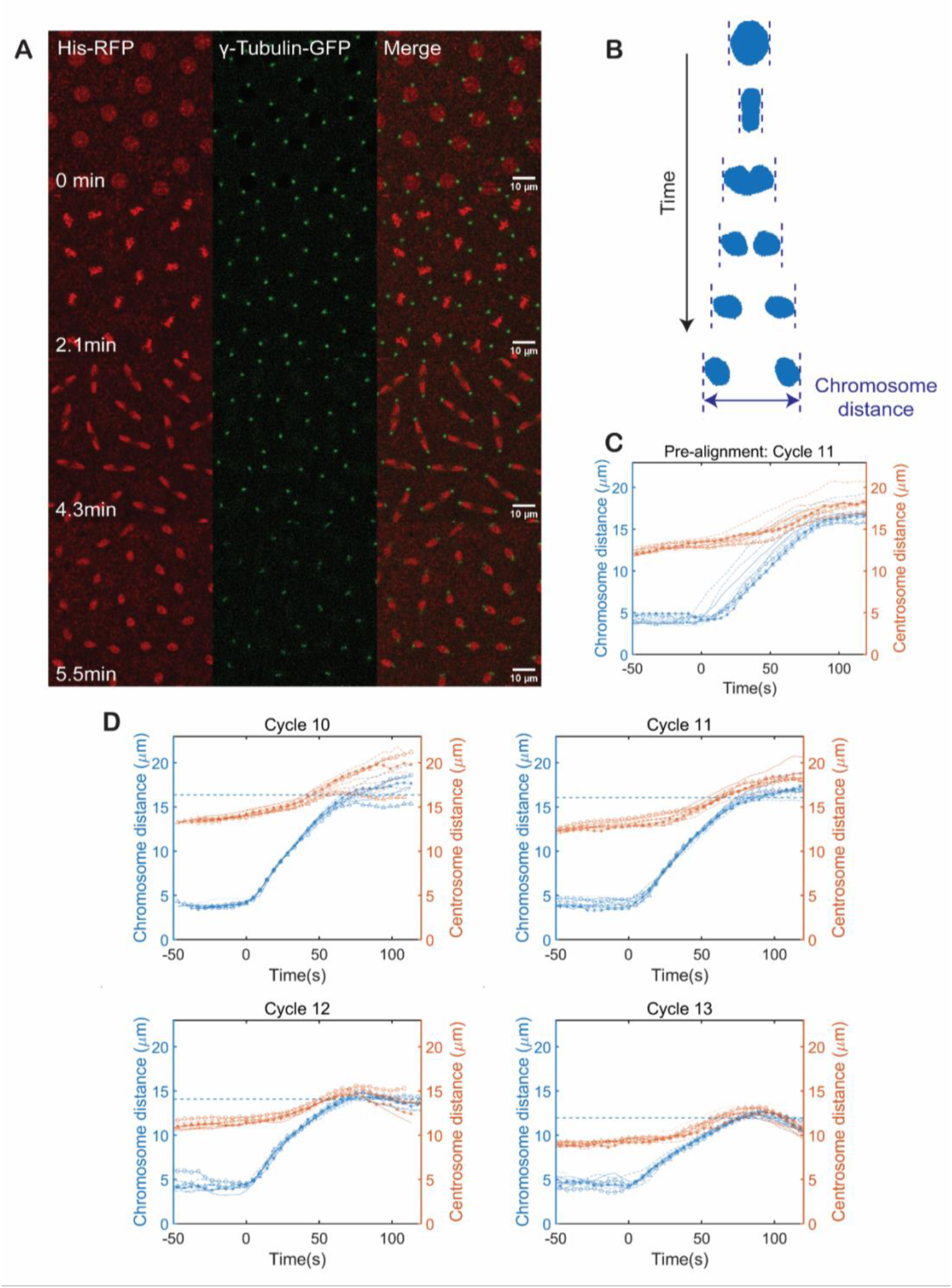
Quantification of chromosome distance, pole-pole distance and approximation of anaphase spindle length. (A) Chromosome movement and centrosome dynamics during mitosis at cycle 11. From top to bottom: prophase, metaphase, anaphase, telophase. Chromosomes are labeled with His-RFP, centrosomes are labeled with γ-Tubulin-GFP. (B) Method for quantifying chromosome distance. Blue masks represent chromosome segmentations. Dashed blue lines indicate the leading edges of chromosomes during separation. (C) Chromosome distance and centrosome distance as a function of time at cycle 11, individual nuclear division events were represented with the same linestyle. A slight asynchrony of chromosome movmement was observed. (D) Aligned chromosome distance and centrosome distance as a function of time from cycle 10 to 13. Chromosome movements were shifted in time to collapase the curve of chromosome distance. T = 0 indicates anaphase onset. Dashed blue line indicates the average of maximum chromosome distance, which serves as a proxy for spindle length in anaphase.

**Figure S2.**
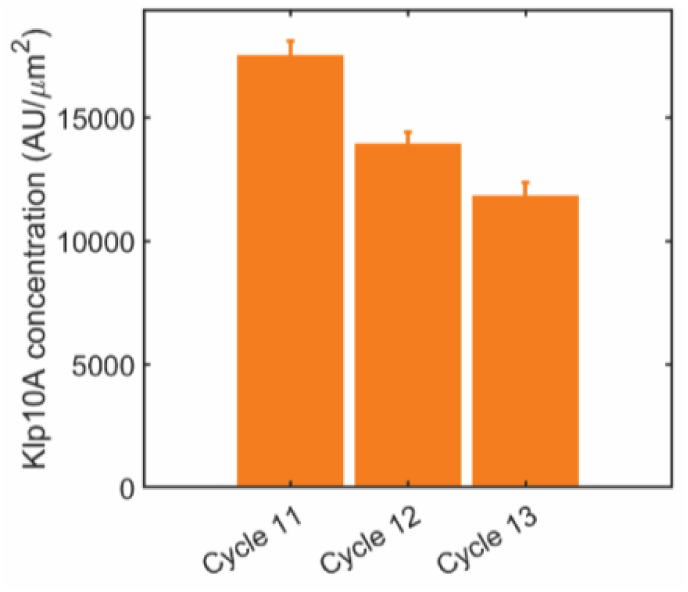
Microtubule depolymerizing motors regulate chromsome velocity. (A) Klp10A concentration at centrosomes is titrated as the cell cycles progress. Data shown as mean ± SEM.

**Figure S3.**
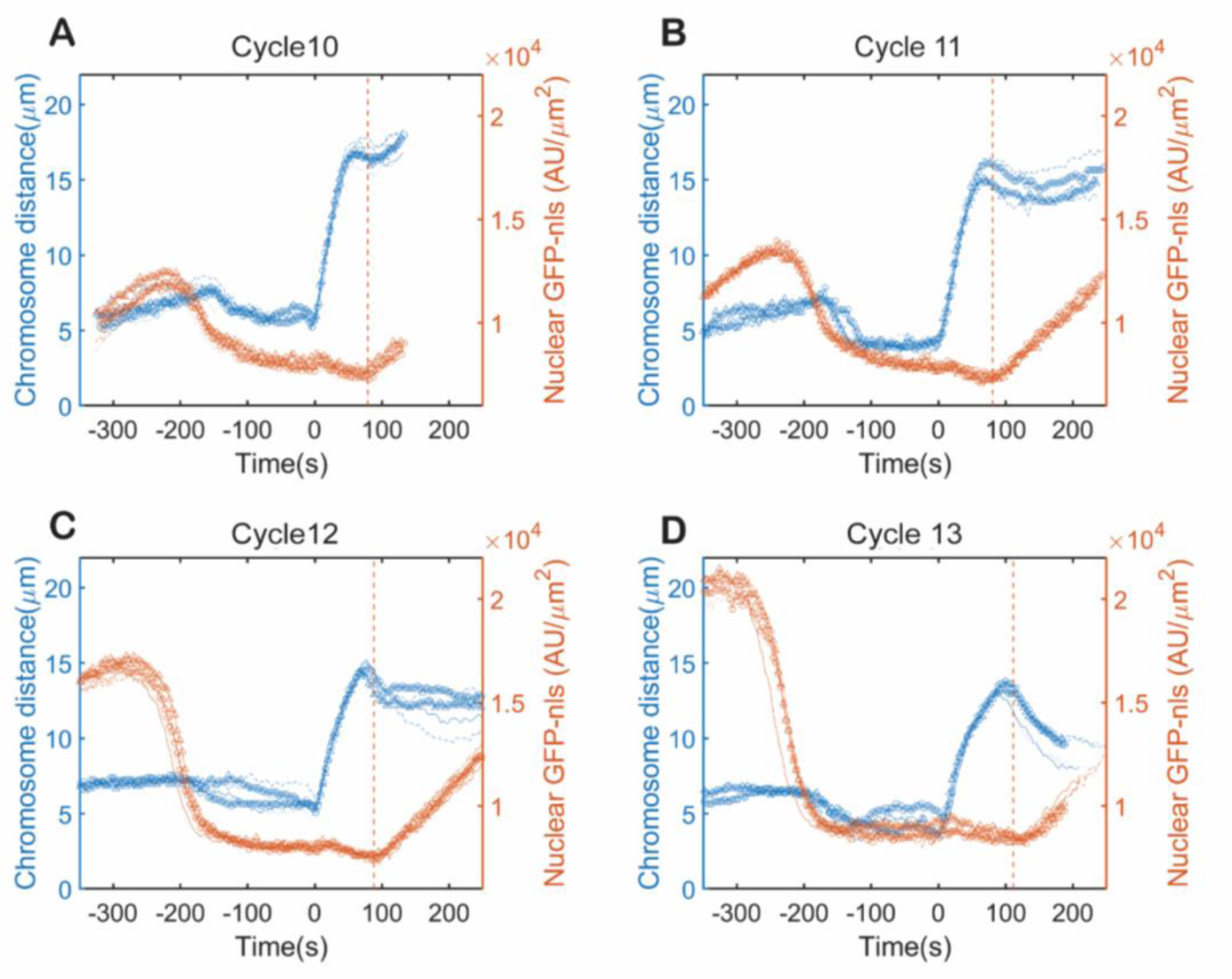
Anaphase duration lengthens as the cell cycles progress. (A-D) Chromosome distance and nuclear localizing GFP concentration as a function of time. T = 0 indicates anaphase onset, dashed orange line indicates the average time point of nuclear envelop reformation.

